# Spread the word about freshwater mussels’ conservation: Increasing awareness and willingness to support conservation-oriented policies through social media

**DOI:** 10.1101/2022.09.09.507251

**Authors:** Noé Ferreira-Rodríguez

**Affiliations:** Universidade de Vigo, Departamento de Ecoloxía e Bioloxía Animal, Vigo, Spain

**Keywords:** digital conservation, netnography, Unionoida, ecological knowledge

## Abstract

Research on social media users has examined the immediate effects of popular social networking sites on vote intention, but studies have not to the potential use of these sites in conservation programs. By conducting an online survey, which included a short-term Before-After-Control-Impact experiment, the author noted an overall increase in respondents’ ecological knowledge after exposure to a dissemination campaign. More importantly, exposure to information through social media affected peoples’ willingness to change their vote intention to support conservation-oriented policies. The current study results suggest that key stakeholders in biodiversity conservation (i.e., researchers, managers, and conservationists) adopting an active role to increase biodiversity valuation may lead to a new form of ecological knowledge acquisition in modern societies.

## Introduction

In recent years, scholars have begun exploring the significance of social media (e.g., Facebook, Twitter, Instagram, Flickr, and Weibo) as effective platforms for fast news dissemination, pluralized voices in reportage, and extensive audience reach in scenarios wherein the spread of information is restricted (Sacco & Bossio, 2015). Moreover, in the case of social conflict, social media has been highlighted as a means for building connections, mobilizing participants and tangible resources, coalition building, and amplifying alternative narratives (Mundt et al., 2018). This emerging field of research reinforces the use of social media networking sites as powerful platforms for reaching wider audiences in modern societies. Social media has been effectively used by politicians to advance their political agendas (Yang et al., 2016). However, few studies have focused on social media in biodiversity conservation (see Toivonen et al., 2019; and references herein).

Despite the increasing political commitment towards biodiversity conservation, significant differences have been noted in emphasis between political priorities and research agendas. Conservation funds are commonly channeled into research priorities determined by politicians, and thus prevent researchers from expanding into new and promising areas (Cairns et al., 2017). In this context, a modern European society must seek to strike a balance between bottom-up (considering citizens as the starting point) and top-down processes (the priorities set by politicians), and integrate it with European policy agendas (Treaty of Lisbon, 2007). Despite responsiveness and participation (i.e., social involvement) serving as the primary values for politicians (Huberts, 2014), researchers’ influence on society—and consequently, in policy-making—has remained largely unexplored (Dassonneville et al., 2020).

In this scenario, the first objective of this study was to explore how key stakeholders in biodiversity conservation (i.e., researchers, managers, and conservationists) can utilize social media to increase people’s ecological knowledge. The second objective was to determine whether increasing people’s ecological knowledge can influence their intention to vote to support conservation-oriented policies. For this purpose, the author explored the utility of social media for fulfilling the outlined objectives in the context of Pearly mussel (Bivalvia, Unionoida; hereafter “mussels”) conservation. Mussels are a diverse and globally distributed group of aquatic organisms found naturally in almost all latitudes, with the exception of Antarctica (Graf & Cummings, 2006). Despite their global distribution, nearly one-third of all species are listed as being threatened with extinction in the International Union for Conservation of Nature Red List (IUCN, 2019), and are thus one of the most endangered animal groups globally. The major causes of their decline are suspected to be caused by increasing demands on global freshwater resources (Downing, 2010). Mussels provide valuable ecosystem services by contributing to biofiltration, nutrient recycling and storage, providing and modifying habitat, and supporting food webs (e.g., Vaughn, 2018). This finding has accelerated research efforts, and contributed toward increasing conservation efforts worldwide (Haag & Williams, 2014). Research on the theoretical basis for protecting freshwater mussels include the identification of key areas for biodiversity conservation, examining propagation and restoration, population genetics, and ecotoxicological studies to document the effects of contaminants on various life stages and derive more stringent water quality criteria and enhanced habitat management (Geist, 2010; Hartmann et al., 2016; Prosser et al., 2017; Strayer et al., 2019; Walters et al., 2019). However, as summarized by Bouska et al. (2018), there are also common challenges in mussel conservation, including lack of public awareness, poor political environment to regulate threats, and hostility toward the management of threatened and endangered species.

By using the social networking site Facebook as an information source, the author constructed a uniquely comprehensive dataset that includes data on ecological knowledge (i.e., knowledge about freshwater mussels’ existence, any contributor to their loss and/or benefit to human well-being) and measures of citizens’ ideological (left–right) positions. The author explored the relationship between citizens’ ideological positions and willingness to act—in this case, through voting for governments that aimed to implement conservation-oriented policies. There are some limitations in this approach. Respondents would be asked about how much they would theoretically sacrifice to preserve an animal group they had not heard about, and whose ecological function they were not aware of (Price, 2000), thus a dissemination campaign was also launched throughout the author’s Facebook page to assess differences between respondents’ willingness to act irrespective of whether they had previous knowledge about this animal group. I hypothesize that providing additional information about freshwater mussels will increase willingness to vote for governments that will implement conservation-oriented policies. With social media becoming one of the main communication channels, the current study results can help managers and conservationists to effectively disseminate information and transform attitudes and behaviors toward the management of threatened and endangered species under conservation programs.

## Material and methods

### Study area

The geographical scope of this study is limited to the autonomous community of Galicia, located in the northwest Spain. Most Galician municipalities are rural and cover 88% of the approximately 30,000 Km^2^ Galician territory, representing 33% of the nearly 2,800,000 Galician population. Although Galicia is moving toward convergence in the expansion of new technologies in territorial information and communication technologies (ICT), considerable differences were noted with other regions in the Information Society in Europe over the past decade (i.e., 32% of the population has never used a computer; Eurostat, 2011).

### Dissemination campaign

Facebook, with 2.6 billion monthly active users, is the most popular social networking site in the world (http://vincos.it/world-map-of-social-networks/). One useful feature of Facebook is the ability to create groups of persons and share or block specific content for each of these groups.

The current study aimed to better understand the potential use of social media for increasing people’s knowledge and valuation of biodiversity, thus the author approached the issue through their own Facebook contacts. Among an original sample of 874 contacts, 400 individuals aged 18 years or older and residing in Galicia were randomly selected for the study. Each potential respondent randomly received a single number, ranging from 1 to 400, and was assigned to the experimental groups. Following a Before-After-Control-Impact (BACI) experimental design, the potential respondents were assigned to the Before-Control group (those who received a number ranging from 1 to 100), Before-Impact group (ranging from 101 to 200), After-Control group (ranging from 201 to 300), or After-Impact group (ranging from 301 to 400).

A dissemination campaign was launched through the researcher’s Facebook page (from February 18 to February 22, 2018; written in Galician). Four posts were developed for the campaign; these posts utilized various elements, including photos, videos, positive statements, and/or questions (Online resource 1), to enhance engagement. The campaign was blind to the After-Control group and included posts on freshwater mussels’ existence, behavior, and host fish dependence, as well as the benefits they provide to human well-being (water purification) and host fish decline by dam construction.

### Social sampling strategy

Following the BACI approach, data was gathered in two rounds via the web by means of a self-administered online questionnaire (Google Forms; written in Galician). Potential respondents were informed about the aim of the study, and voluntary participation, anonymous response and confidentiality. After informed consent was obtained, the link to the questionnaire was disseminated through a private chat. The first round (before the dissemination campaign) was disseminated to the Before-Control and Before-Impact groups on February 15. The second round (after the dissemination campaign) was disseminated to the After-Control and After-Impact groups on February 22. The questionnaire was closed on March 22, when no more answers were received. Respondents were allowed to share the questionnaire (link to the questionnaire) with others on their social media profiles.

### Online questionnaire

The questionnaire was designed to investigate the following aspects: (1) general knowledge about freshwater mussels, (2) the potential use of social networks in improving ecological knowledge and biodiversity valuation, and (3) how increasing people’s ecological knowledge and biodiversity valuation can influence their vote intention.

Potential respondents were asked to participate in the study and respond to a questionnaire related to their knowledge about freshwater mussels. Respondents were informed that all responses were anonymous, and there were no correct or wrong answers. The questionnaire included an open-ended question about overall knowledge on freshwater mussels, major contributors to freshwater mussels’ loss, and benefits they provide for human well-being, followed by statements in which respondents selected four options from a list of major contributors to loss and benefits to human well-being. A “do not know” category was included for each question. Respondents were inquired about their willingness to vote for a different political party—other than their usual choice—depending on political commitment to freshwater mussels’ conservation. To ensure that the sampled population fairly represents the distribution within the left–right political spectrum, the questionnaire also inquired about their political ideology. Finally, respondents were asked about socio-demographic characteristics (i.e., age, income, education level, and place of residence and origin; Online resource 1, English version).

### Statistical analysis

Descriptive analysis was used to estimate the percentage of respondents with any knowledge about freshwater mussels, including major contributors to their loss and benefits they provide to human well-being. An ordinal measure of the social importance of each contributor and benefit was created based on the number of times a given answer was reported. Ordinal logit regression (logit) was used for predicting respondents’ ecological knowledge about freshwater mussels. To compare respondents’ willingness to vote for a different political party based on their political commitment to freshwater mussels’ conservation, ordinal logistic regression was used to test relationships, depending on their position on the left–right political spectrum. To assess changes in users’ knowledge about freshwater mussels and willingness to change vote intention, the post-impact group was compared against the control group. To ensure these changes are related to the dissemination campaign and not to other sources of external information, the control group was also compared against the pre-impact group. The answers were also linked to the socio-demographic information (i.e., age, income, education level, and place of origin) of respondents. Statistical tests were conducted using the *glm* package in R (version 3.3.1).

## Results

A total of 110 respondents (i.e., 27.5% response rate) completed the interview. There were 21 respondents in the Before-Control group, 28 respondents in the Before-Impact group, 41 respondents in the After-Control group, and 20 respondents in the After-Impact group.

Overall, half of respondents had prior knowledge about freshwater mussels. However, significant differences were noted before and after the dissemination campaign (p<0.05). Specifically, for one individual who was exposed to the dissemination campaign, the odds of being aware of freshwater mussels’ existence (versus being in the control group) increased by 0.18.

Among respondents with prior knowledge about freshwater mussels’ existence, most (61%) were aware of any contributor to the loss of freshwater mussels, whereas 39% were unaware. Most respondents considered the loss of freshwater mussels to be related to pollution, alien invasive species, and flow alteration by dam construction. In contrast, land-use change, predation, and collection were rarely perceived as relevant causes of freshwater mussels’ loss (Figure 1a). Most respondents with prior knowledge about freshwater mussels (78%) also perceived that they provide benefits contributing to the population’s well-being, whereas less than 11% considered that freshwater mussels do not provide any benefits. Regarding the perceived importance of ecosystem services, freshwater mussels as food for other animals, water quality and water purification were considered as the most important factors. In contrast, their utilization as ornaments and cultural value were rarely identified as relevant ecosystem services (Figure 1b). The dissemination campaign had no effect at any significance level in increasing respondents’ knowledge about contributors to their loss or benefits for human well-being.

**Figure 1.**
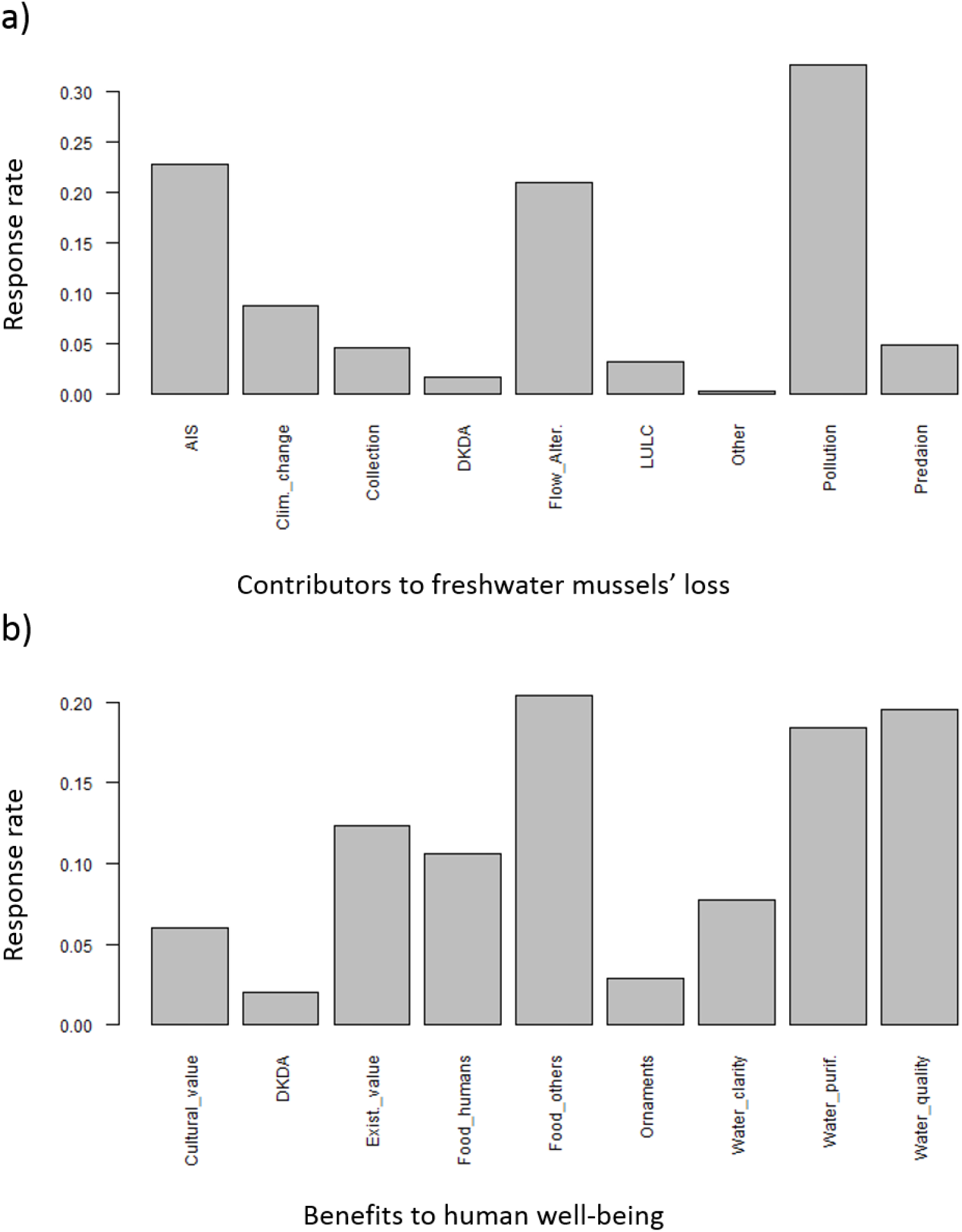
General knowledge about freshwater mussels. (a) Perceived contributors to freshwater mussels’ loss, and (b) social importance perceived for ecosystem services (AIS: Alien invasive species; DKDA: Do not know or did not answer; LULC: Land use and land cover change).

Being on the left side of the political spectrum and selecting other political preference, versus being in the center of the left–right political spectrum, changes the log odds of being aware about freshwater mussels by 1.63 and 2.11, respectively. For a one-unit increase, for one individual on the left side of the political spectrum, the odds of being aware of freshwater mussels’ existence increased by 5.12. Similarly, selecting other political preference increased the odds of being aware about freshwater mussels by 8.25 (Table 1).

**Table 1.** Results of logistic regression models of being aware about freshwater mussels’ existence for people in the left–right political spectrum.

| Political spectrum | Estimate | Std. Error | z value | p | Odds ratio<sup>a</sup> (and 95 % CL) |
| --- | --- | --- | --- | --- | --- |
| Intercept | −1.504 | 0.782 | −1.924 | 0.054 | −0.222 (0.034–0.862) |
| Left | 1.633 | 0.822 | 1.987 | 0.047* | 5.120 (1.199–35.423) |
| Right | 0.811 | 1.453 | 0.558 | 0.577 | 2.250 (0.081–40.328) |
| Other | 2.110 | 0.932 | 2.264 | 0.024* | 8.250 (1.525–67.179) |

Signif. codes: 0 '\*\*\*' 0.001 '\*\*' 0.01 '\*' 0.05 '.' 0.1
<sup>a</sup> Odds of having knowledge about freshwater mussels' existence.

Willingness to vote for a different party before and after the dissemination campaign was marginally significant (p<0.1, Table 1). Specifically, being exposed to the dissemination campaign, versus being in the control group, changes the log odds of voting for a different party by 1.39. For a one-unit increase, for an individual exposed to the dissemination campaign, the odds of voting for a different party (versus voting for the same party as they usually do) increased by 4 (Table 2).

**Table 2.**
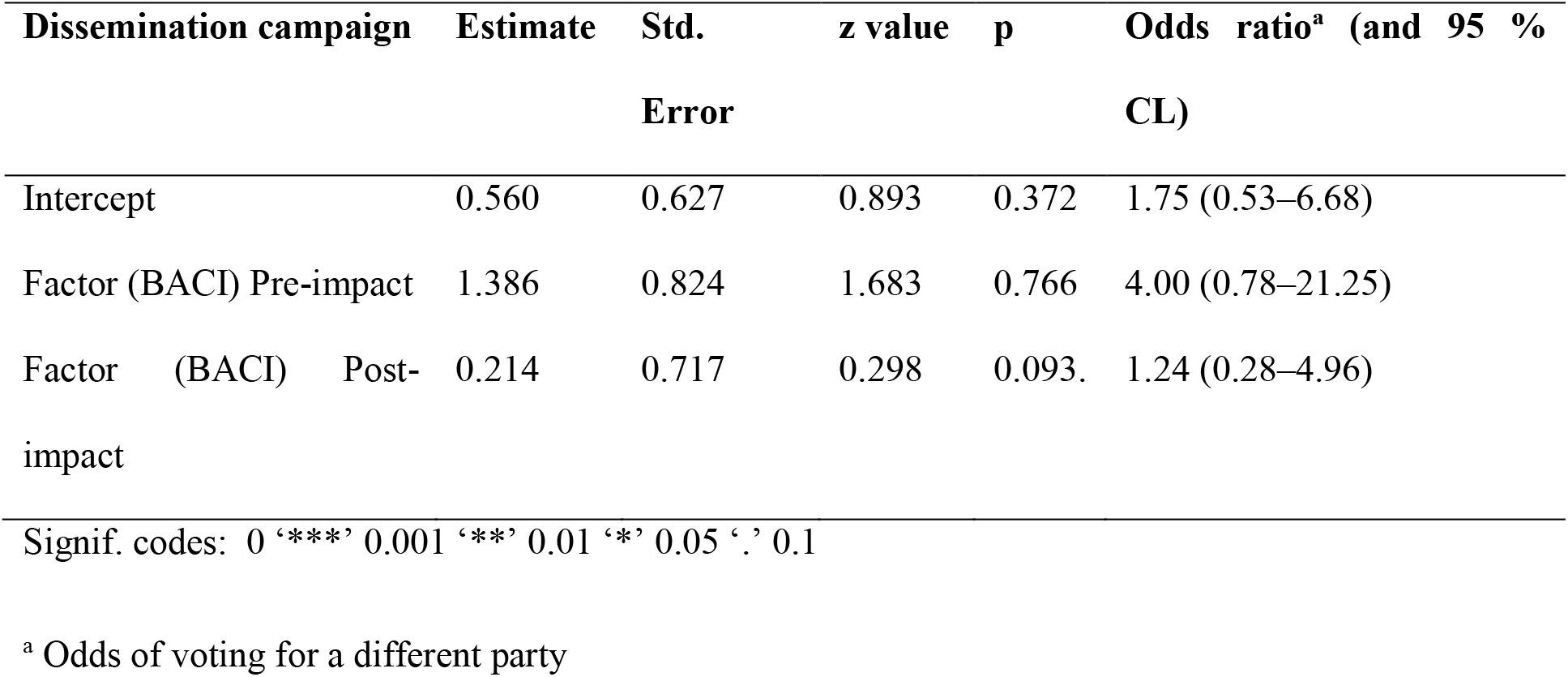
Results of logistic regression models for people’s willingness to vote for a different party before and after the dissemination campaign

## Discussion

Research on social media users has examined the immediate effects of popular networking sites on vote intention, but studies have yet to examine its potential use by key stakeholders (e.g., researchers, managers, and conservationists) in conservation programs. The primary objectives of this study were to explore the utilization of social media for increasing people’s ecological knowledge and biodiversity valuation, and determining whether increasing people’s ecological knowledge and biodiversity valuation can influence their vote intention to support conservation-oriented policies. The results show that dissemination through social media increased people’s ecological knowledge and, in particular, knowledge about freshwater mussels. Results also show that increasing people’s ecological knowledge can affect peoples’ willingness to change their vote intention. Therefore, the current study is important because lack of public awareness and poor political environment for regulating threats are common challenges to be addressed in mussel conservation programs.

The fundamental processes related to mussels’ abundance and behavior are of considerable relevance for regulating ecosystem services in freshwater environments (Oosterhuis et al., 2021). However, previous information received by the population for acquiring this ecological knowledge was either non-existent or disseminated through non-effective channels. As a consequence, ecological knowledge about freshwater mussels’ existence, contributing factors to their decline, and the benefits they provide, remain largely unknown by the population. The probability of one individual becoming aware about the existence of freshwater mussels was, in addition, related with the political preferences of the sampled population. Nevertheless, results have shown that social media can be an effective channel for increasing ecological knowledge and their influence may affect public willingness to vote for governments that would implement conservation-oriented policies.

From the sampled population, only half of them were aware about freshwater mussels’ existence and less were aware about the threats they face or benefits they provide. However, some respondents were willing to change their vote depending on political commitment to freshwater mussels’ conservation, although they did not know freshwater mussels exist (i.e., 28 respondents out of 54). Hence, it is assumed that freshwater mussels attracted a symbolic behavior for conservation, which is reflected as a sense of wellbeing, of simply knowing freshwater mussels exists, even if they will be never utilized or experienced. Those who are willing to change their political vote should be considered as people concerned about nature conservation or, according to Price (2000), as people “expressing desire to be [seen to be] acting responsibly towards nature.” It should be noted, however, that willingness to change political vote significantly increased among those who received extra information. Hence, we can assume that key players in biodiversity conservation can raise public awareness toward biodiversity conservation and, in consequence, could reorient conservation in more ethically expansive directions that incorporate recognition of the intrinsic value of wildlife (Wallach et al., 2018). The question is, however, whether this public awareness created through social media campaigns may encourage the implementation of conservation-oriented policies. In this regard, studies have shown that politicians are responsive to the opinions of voters, but not to those of non-voters (Dassonneville et al., 2020). Hence, it is increasingly important to assess whether there is an empirical nexus between knowledge transference, voters’ choice, and conservation policies.

The costs associated to online information dissemination are related to the rapid spread of misinformation and public access to large amounts of knowledge from unconventional classes of experts. However, it also represents a unique opportunity to learn about biodiversity related issues and the benefits of online information dissemination significantly outweigh the costs associated with internet cacophony (Pierpoint, 2011). On the other side, although the benefits of online surveys are their ease-of-access for respondents, and savings in time and economic resources (Wright, 2005), there are several limitations. The approach used in this study (allowing people to share the questionnaire link in their profiles) may have introduced some bias in the study because it was possible that (1) some respondents did not fulfill the criteria for being included in the study (e.g., individuals aged 18 years or older residing in Galicia) or people answering the survey who received the link from someone other than the researcher were not correctly assigned to the right group. The questionnaire itself was spread via Facebook, which may have excluded people in younger and older age groups. In terms of age, majority of the respondents (60.92%) were aged between 30 and 40, whereas the second more prominent (15.45%) age group involved people aged between 40 and 50. To visualize the sample distribution within the left-right political spectrum, the sample was compared with national information of political preferences in Spain, which were conducted by the Sociological Research Center (CIS, 2017). According to official data, 37.7% of the population is positioned on the left or on the center-left side of the political spectrum, 31% on the center, and 15.1% on the center-right or right. Based on this information, the sample is biased toward respondents positioned on the left side of the political spectrum. In addition, according to official data, respondents selecting other political preferences or not answering the question are likely to be positioned on the center or on the right side of the political spectrum. Finally, the low response rate (*ca*. 25%) could be related with voluntary response bias because individuals who chose to respond to the questionnaire were self-selected and may have had more knowledge about freshwater mussels, compared with those who chose not to respond. Refusal to answer (or registration of zero or infinite willingness) was termed as protest responses (Price, 2000). Altogether, this information may provide valuable information for designing future dissemination campaigns in biodiversity conservation programs by using social media platforms to increase public awareness and willingness to support conservation policies.

## Acknowledgements

I thank all the people who generously responded to my interview. I am grateful to the two anonymous reviewers and the (associate) editor of Human Dimensions of Wildlife for their valuable comments and suggestions. The work of the author was supported by a postdoctoral fellowship from the government of the autonomous community of Galicia (Xunta de Galicia, Nº de exp. ED481D-2021-023).

## Notes

Local ethical approval was not acquired in 2018 as there was no Ethical Committee for social science research at the university (Universidade de Vigo) at that time.

